# Multi-Modal Disentanglement of Spatial Transcriptomics and Histopathology Imaging

**DOI:** 10.1101/2025.02.19.638201

**Authors:** Hassaan Maan, Zongliang Ji, Elliot Sicheri, Tiak Ju Tan, Alina Selega, Ricardo Gonzalez, Rahul G. Krishnan, Bo Wang, Kieran R. Campbell

## Abstract

Spatially-resolved expression profiling data has revolutionized biological research with multiple emerging clinical applications. Spatial transcriptomic assays are often jointly measured with histopathology imaging data, which is frequently used for diagnosing and staging various diseases. However, determining the extent to which the spatial transcriptomic and histopathology data represent overlapping or unique sources of variation is challenging, particularly given the myriad of factors influencing both, including expression variation, spatial context, tissue morphology, and batch effects. Here, we view this challenge as multi-modal disentanglement and develop an evaluation framework. We introduce SpatialDIVA, a disentanglement technique for jointly measured spatially resolved transcriptomics and histopathology data. We demonstrate that SpatialDIVA outperforms baseline techniques in disentangling salient factors of variation in curated pathologist-annotated multi-sample colorectal and pancreatic cancer cohorts. Further, SpatialDIVA removes batch effects from multi-modal data, allows for factor covariance analysis, and yields actionable biological insights through a novel conditional multi-modal generation method. The SpatialDIVA model, evaluation code, and datasets are available at https://github.com/hsmaan/SpatialDIVA.

## 1 Introduction

Spatial context is an important measurement in the study of biological systems, as it dictates the organization of tissues, the flow of information in the form of cell to cell communication, transport of biomolecules and nutrients, and many other factors (Rao et al., 2021; Tian et al., 2023). To incorporate spatial context in molecular measurements, researchers have developed spatial transcriptomics (ST) technologies, which quantify mRNA expression in small numbers of cells (1-10) at specific locations in a tissue, commonly referred to as *spots* (Moses & Pachter, 2022). Barcoding then allows for identifying the spatial position of captured RNA molecules (Moses & Pachter, 2022). After determining cell-type identity through the transcriptomic information, the spatial context can be used to perform additional analysis, such as annotating spatial domains and determining patterns of cell to cell communication (Rao et al., 2021).

In addition to quantifying spatially-resolved gene expression, spot-based ST assays such as 10X Visium (Ståhl et al., 2016) are able to simultaneously image a hematoxylin and eosin (H&E) stain (Fischer et al., 2008; Janesick et al., 2023) of the tissue section under study for histological analysis. H&E staining of tissue and examination of associated morphological features is routinely used for diagnosing and staging many malignancies (He et al., 2012). The addition of H&E staining to ST technologies allows researchers to associate morphological features with RNA expression of different genes in the underlying tissue, as well as predict gene expression from H&E (Janesick et al., 2023).

Although there is a myriad of existing approaches to predict RNA expression from histopathology images (Sec. 5), these are limited, as histopathology data is often only weakly correlated with RNA expression (Zeng et al., 2022; Xie et al., 2023). Further, this correlation is highly variable across genes (Zeng et al., 2022; Xie et al., 2023). One way to interpret these methods is that they are approximating the upper bound on mutual information between the two modalities. However, this upper bound does not answer the question of *which information* is present or absent in each modality, and this is more biologically relevant. For example, knowing which variation is exclusively found in RNA expression and not in histology data allows us to determine which factors relevant to disease states are being missed through only performing histology analysis, and vice versa.

Many other factors also contribute to modality-specific variation, such as spatial context, and nuisance factors including batch effects. Delineating the contribution of these factors allows us to obtain an interpretable model of the multi-modal data generating distribution, infer which gene-programs and morphological patterns are dictated by specific factors, and remove the effects of nuisance variation.

In this work, we determine the precise effects of different biological and technical factors in multi-modal ST and histology data. Our contributions are as follows:

- In Sec. 2.1-2.3, we frame the challenge of determining the contribution of distinct biological factors to multi-modal assays as a multi-modal disentanglement problem. We propose a general framework that incorporates biological knowledge (Appendix A.1), based on prior work that demonstrates fully unsupervised approaches to disentanglement are suboptimal (Locatello et al., 2018).
- We introduce *SpatialDIVA* (Fig. 1) in Sec. 3, which builds upon previous work in disentangled representation learning (Ilse et al., 2019) by introducing prior-constrained multi-modal disentanglement in a ST and histopathology setting, with continuous label distributions.
- We show that SpatialDIVA outperforms previous techniques and exhibits strong disentanglement of colorectal and pancreatic cancer data (Sec. 4.1 and 4.2). To benchmark disentanglement in this setting, we develop a framework incorporating several metrics and two multi-patient/sample pathologist-annotated datasets, which can be used by the community to further our contributions (Appendices C and H).
- To determine modality-specific effects of latent factors, we develop a conditional generation framework for multi-modal data using the SpatialDIVA model (Fig. 1), and use this framework to infer modality-specific gene regulatory programs in a multi-patient pancreatic cancer cohort (Sec. 4.3 and 4.4).

**Figure 1:**
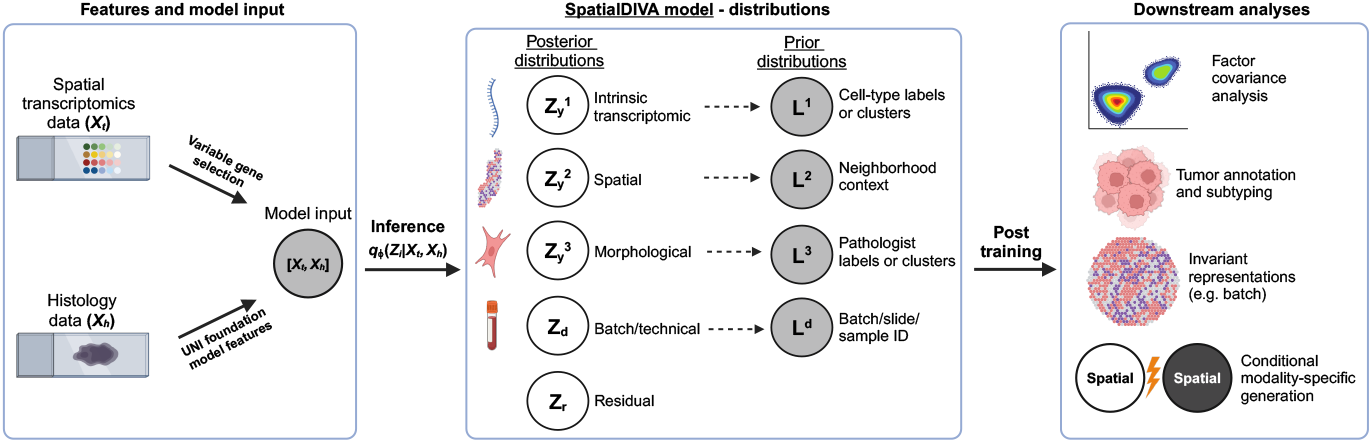
Overview of contributions and the *SpatialDIVA* model. We frame the challenge of evaluating modality and factor-specific contributions as a multi-modal disentanglement problem, for which we present the SpatialDIVA model as a solution. SpatialDIVA allows for biologically and clinically relevant downstream analyses, such as factor covariance, tumor annotation in cancer biopsy samples, removing batch effects, and determining modality-specific information through conditional generation.

## 2 Background and problem formulation

### 2.1 ST and histology data

Given a slide with a slice of tissue and jointly measured ST and histology (Ståhl et al., 2016), we obtain the following data representation per spot *i* on the slide:

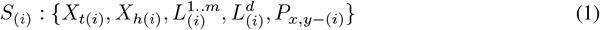

Where *X*_t_ indicates the transcriptomic counts across genes measured in the spot, *X*_h_ indicates the histopathology image distribution, *L*^1..m^ indicates any *m* label groups for the spot (such as pathologist annotations and cell-type labels), *L*^d^ indicates a label for the specific slide or tissue section (e.g. slide 1, slide 2, …), and *P*_x,y_ are the spatial coordinates of that spot on the slide.

The distribution of expression of each gene *j* per spot *i* is given by *g*_*i,j*_ ∼ Poisson(*λ*_*i,j*_). Log-normalization of this count data can be performed for modelling (Hao et al., 2024), or the untransformed counts can be used directly (Zhao et al., 2022) (Appendices D and E). The distribution of the histopathology data follows a 3-channel RGB image, and histopathology foundation models can be used to extract features for this data (Chen et al., 2024) (Appendix D).

### 2.2 Latent variable model of ST and histology

The underlying biology that drives the variation of ST and the paired histology data is often shared. However, the different biological and technical factors, and the extent to which they contribute to each modality, is unresolved.

Assume a set of *k* generative factors *V* = {*v*_1_, *v*_2_, …, *v*_*k*_} account for the data distributions of an arbitrary number of views 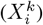 through an arbitrary number of generative processes *G*(.)^k^:

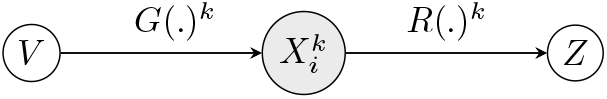

The underlying generative processes *G*(.)^k^, as well as the generative factors *V*, are unobservable. For an arbitrary number of observed views, our goal is to approximate the generative factors jointly for all views through processes *R*(.)^k^ and infer *m* latent variables (*Z* = *z*_1_, *z*_2_, …, *z*_m_):

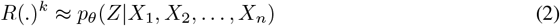

In the context of spatial transcriptomics, we know that both ST (**X**_**t**_) and histology (**X**_**h**_) are generated by overlapping biological factors, which we can approximate through *m* latent variables and the *R*(.)^t^ and *R*(.)^h^ processes :

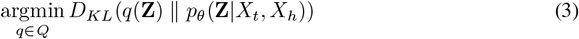

Note that any number of views can be considered in this framework, but we restrict it to two based on our joint ST and histology setting. We aim to learn parameterizations for the functions *R*(.)^t^ and *R*(.)^h^. Assuming that these functions are parameterized by *ψ*_1_ and *ψ*_2_, collectively described as *ψ*, the objective becomes:

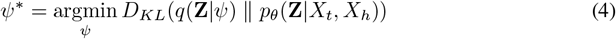

### 2.3 Disentangling explanatory factors for ST and histology

Given that we want to learn *m* explanatory factors for both the ST (**X**_**t**_) and histology (**X**_**h**_) data that best approximate the underlying generative distribution corresponding to distinct biological processes, we aim to learn *disentangled* representations of the approximated latent distribution *Z*. In general, we want to minimize the total correlation between each learned latent covariate:

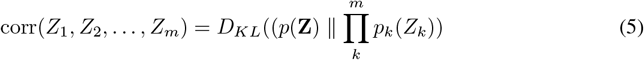

The other constraint in our setting is that we want the learned latent factors to correspond to both known and novel biological sources of variation.

It has been shown by Locatello et al. (2018) that unsupervised approaches to learning disentangled representations, such as through the Beta variational autoencoder (*β*-VAE) model (Higgins et al., 2016), are under-defined, and entangled representations can lead to the same marginal distributions as disentangled representations in this setting. This renders unsupervised identification of non-linear latent variables difficult. Therefore, we incorporate relevant prior biological knowledge (Appendix A.1). Further, we add a residual latent factor that accounts for any variation not captured by prior-constrained factors, similar to previous work (Ilse et al., 2019).

## 3 The SpatialDIVA model

Given the problem description outlined in Sec. 2, we introduce a generative framework for disentangling ST and histology data: the Spatial Domain Invariant Variational Autoencoder (SpatialDIVA) model (Fig. 1 and 2). SpatialDIVA is a deep latent variable model, that aims to infer latent distributions for *m* biological covariates 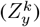, batch effects (*Z*_d_), and residual variation (*Z*_r_), through maximizing the marginal likelihood of the histology (*X*_h_) and ST (*X*_t_) data, as well as prior knowledge (*L*):

**Figure 2:**
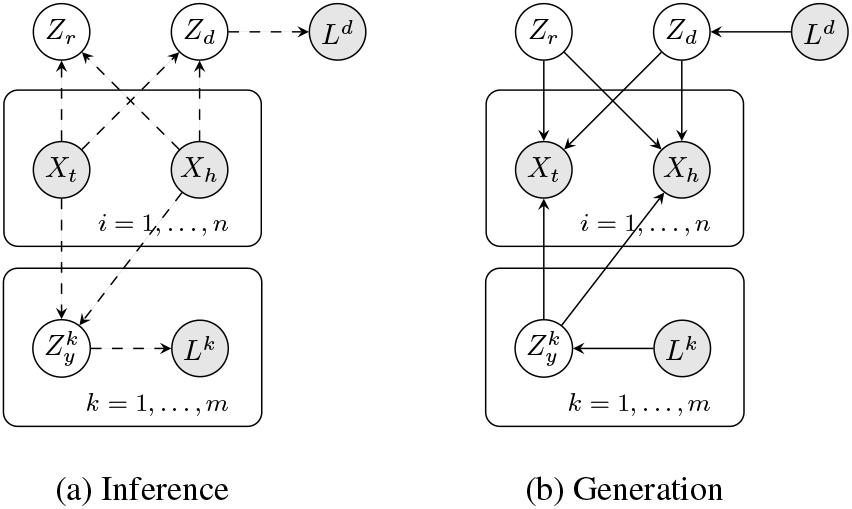
SpatialDIVA model overview. **(a)** The observed ST (*X*_t_) and histology (*X*_h_) representations are used to infer latent residual variation (*Z*_r_), batch/technical variation (*Z*_d_), and variation of key biological factors 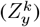, including intrinsic transcriptomic, morphological, and spatial factors. **(b)** The *X*_t_ and *X*_h_ likelihoods are generated by conditioning on learned factors, where the conditioning is controllable.

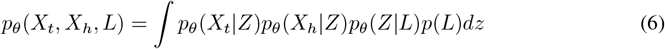

However, this marginalization is intractable, and therefore we learn a lower bound on the log-likelihood (Appendix F):

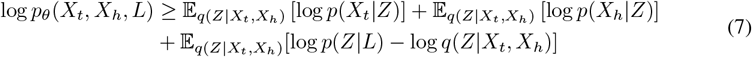

Incorporating SpatialDIVA’s prior-constrained latent variables (Appendices A.1 and F), we obtain our evidence lower bound (ELBO), where *θ* and *ϕ* are neural network parameters (Appendix E):

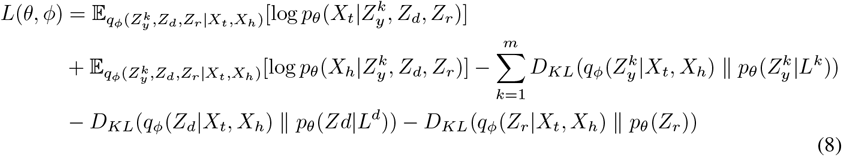

Similar to DIVA (Ilse et al., 2019) and CCVAE (Joy et al., 2020), we incorporate classification heads parameterized by *ψ*, to ensure that the posterior distributions contain the relevant labeled knowledge (*L*), giving the overall objective:

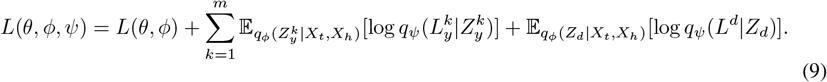

We encode the **batch labels (***L*^d^) using an index of the slide from which each spot originates, enabling generalization across slide contexts. **Prior biological knowledge** is encoded based on labels that contain biologically relevant information from both modalities (*L*^k^) (Appendices A.1 and G).

For **intrinsic transcriptomic variation**, we use a categorical distribution of expert-annotated cell-type labels for each spot (Appendix G). For **morphology variation**, we use pathologist annotations of morphology features on histology slides corresponding to each spot, through a categorical distribution (Appendix G). In cases where prior annotations might not be available, unsupervised clustering on the ST and histology data individually can be used to derive modality-specific labels.

The last biologically informative prior that we consider is **spatial context**. Each spot on a spatial transcriptomics slide has spatial coordinates *P*_x_, *P*_y_. However, these coordinates are not generalizable across slides. Therefore, we developed a **spatially aware context distribution** for each spot (Appendix G). For each spot *i*, we determine the *N* nearest neighbors using the spatial coordinates (*P*_x_, *P*_y_). For both the transcriptomic (*X*_t_) and histology (*X*_h_) features, we decompose their information for all spots in a slide through principal component analysis (PCA):

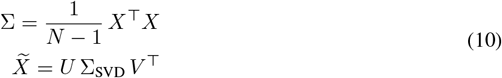

We use the concatenated decomposed representations of *X*_t_ and 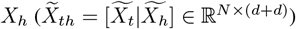 of the *N* neighbors for spot *i* to obtain a representation of spatial context (*Y* ^s^):

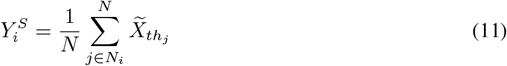

A key advantage of this approach is that the model does not have to predict the high-dimensional ST and histology distributions, leading to faster training when used in the classification and prior distribution losses (Eqn. 9) (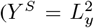 for spatial context). This generalizable representation of spatial context can be used across slides/samples, and allows for multi-sample training. Complete details on all of the biological priors can be found in Appendices A.1 and G.

Within this problem context, SpatialDIVA offers significant advantages compared to other approaches for modelling multi-modal ST and histology data (Table 1, Sec. 5).

**Table 1:** SpatialDIVA compared to other methods for jointly modeling ST and H&E image data.

| Method | Joint embeddings? | Disentangled representations? | Interpretable latent space? | Conditional multi-modal generation? |
| --- | --- | --- | --- | --- |
| **SpatialDIVA** | ✓ | ✓ | ✓ | ✓ |
| Unsupervised disentanglement | ✓ | ✓ | ✗ | ✗ |
| Histology to gene expression | ✓ | ✗ | ✗ | ✗ |

## 4 Results

### 4.1 Evaluating disentanglement of multi-modal ST and histology data

To assess how well SpatialDIVA can disentangle factors of variation affecting both modalities, we compared it with baseline models using a series of disentanglement metrics across pathologist-annotated datasets.

#### Datasets

We collated multi-patient pancreatic ductal adenocarcinoma (PDAC, 13 slides) and colorectal cancer (CRC, 14 slides) data profiled with ST and H&E imaging (Cui Zhou et al., 2022; Valdeolivas et al., 2024). We created pathologist annotations for tumour/normal tissue regions for the PDAC dataset and used existing pathologist annotations for the CRC dataset (Appendix C). We preprocess the data such that we obtain patches of the H&E image that correspond to the area around the spots that capture ST data (Appendix C). Image features for a per-spot representation are obtained through zero-shot inference of the UNI histopathology foundation model (Chen et al., 2024). ST data is processed uniformly for all datasets (Appendix C), and highly-variable gene (HVG) selection is performed for all spots across all slides (Appendix D).

#### Baselines

We compared SpatialDIVA to an array of baselines including PCA, a standard VAE model (Kingma & Welling, 2013), and an unsupervised disentanglement approach in the *β*-VAE model (Higgins et al., 2016) (Appendix H). Comparison with the DIVA model (Ilse et al., 2019) and similar approaches (Joy et al., 2020) was not possible as they cannot handle multi-modal data and continuous label distributions for spatial context (Appendix E).

#### Evaluation

A disentangled representation would result in each factor containing specific information about morphology, transcriptomic, spatial, and batch variation. For morphology we used pathologist annotations of the data at a per-patch level. For transcriptomic information, we used expert-annotated cell-types within each spot. For batch variation, we used the slide label that each spot originates from. It was unclear how to evaluate continuous spatial variation in this case, so this was omitted. The categorical labels for the morphological, transcriptomic, and batch variation were one-hot encoded and embeddings from the baselines and SpatialDIVA model were compared with the encoded labels to assess disentanglement (Appendix H).

Models were trained on randomly selected 90% subsets of the datasets for PDAC and CRC (one model per cancer type), then evaluated on the held-out 10% of the data for 10 iterations (Appendix H). Disentanglement was assessed using multiple metrics for categorical factors and continuous embeddings (Appendix I) (Carbonneau et al., 2020).

Overall, SpatialDIVA performed the best considering an aggregate ranking across all disentanglement metrics used in the benchmark, for both the PDAC and CRC cohorts (Table 2). These results demonstrate that within a multi-modal disentanglement setting, the conclusions from Locatello et al. (2018) hold, and that supervision based on prior information leads to better disentanglement.

**Table 2:** Disentanglement benchmark results for the pancreatic (PDAC) and colorectal cancer (CRC) datasets. Results are the mean scores across 10 random subsamples of the datasets. The best results for each metric are in **bold** and the second best results are <u>underlined</u>. Average rank is based on rankings across all metrics for each dataset (Appendix H). Standard deviations across iterations can be found in Appendix B (Table 4).

| Method \ Metric | Colorectal cancer cohort |  |  |  | Pancreatic cancer cohort |  |  |  |
| --- | --- | --- | --- | --- | --- | --- | --- | --- |
| | PCA | VAE | β-VAE | SpatialDIVA | PCA | VAE | β-VAE | SpatialDIVA |
| JEMMING (↑) | 0.2011 | <u>0.7690</u> | **0.9200** | 0.3537 | 0.2579 | <u>0.7922</u> | **0.9032** | 0.5327 |
| SAP (↑) | **0.0049** | 0.0000 | 0.0000 | <u>0.0024</u> | **0.0203** | 0.0000 | 0.0000 | <u>0.0034</u> |
| MIG (↑) | <u>0.0075</u> | 0.0049 | 0.0041 | **0.0128** | <u>0.0225</u> | 0.0057 | 0.0038 | **0.0893** |
| MIG-SUP (↑) | 0.0102 | **0.0541** | 0.0328 | <u>0.0336</u> | 0.0186 | **0.0668** | 0.0246 | <u>0.0577</u> |
| Modularity (↑) | 0.9473 | <u>0.9600</u> | 0.9562 | **0.9647** | **0.9421** | 0.9298 | 0.8938 | <u>0.9357</u> |
| DCI-MIG (↑) | <u>0.0337</u> | 0.0007 | 0.0003 | **0.1231** | <u>0.0700</u> | 0.0001 | 0.0001 | **0.2131** |
| Explicitness (↑) | **0.9505** | 0.1032 | 0.0409 | <u>0.9375</u> | <u>0.9010</u> | 0.0293 | 0.0035 | **0.9433** |
| IRS (↑) | 0.5008 | **0.5196** | 0.5001 | <u>0.5085</u> | 0.4459 | 0.4951 | <u>0.5000</u> | **0.5351** |
| Average Rank (↓) | 3 | <u>2</u> | 4 | **1** | <u>2</u> | 3 | 4 | **1** |

### 4.2 Assessing disentanglement and covariance of latent factors

To determine how the disentanglement properties of SpatialDIVA affect the learned latent spaces, we trained the model on the 13 PDAC slides and evaluated samples from the posterior distributions 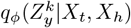 and *q*_ϕ_(*Z*_d_ |*X*_t_, *X*_h_), conditioned on the observed data *X*_t_ and *X*_h_ (Appendix H). We performed PCA on high-dimensional samples from one factor and visualized the first two PCs. We then overlaid the label distributions onto the PCA reduction of each learned factor, to visually examine how the annotated labels covary in the factor-specific latent spaces (Fig. 3 and 4).

**Figure 3:**
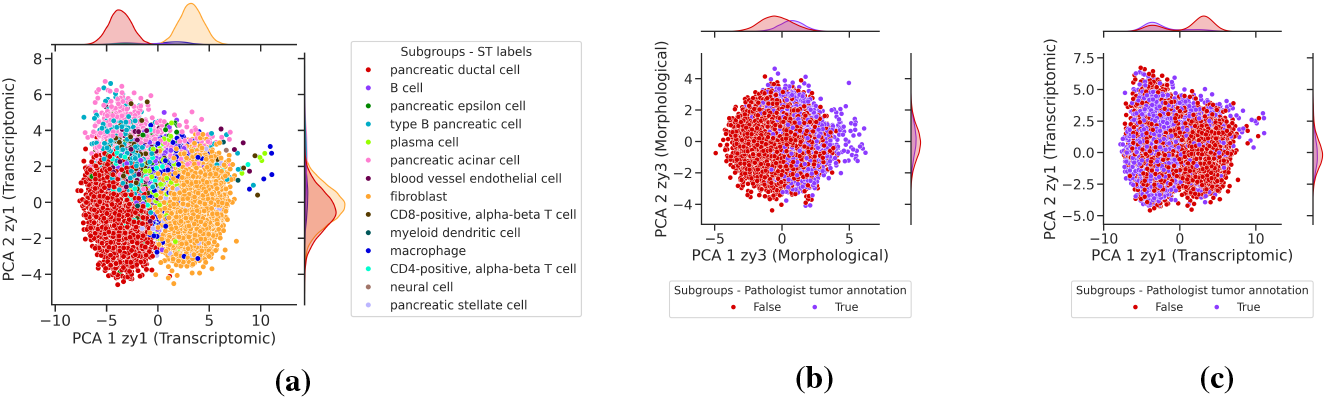
Transcriptomic and morphology-associated posterior distributions. High-dimensional samples from the posterior distributions 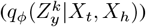 of the transcriptomic-associated (a, c) and morphology-associated (b) latent spaces, reduced using PCA and the two axes associated with the highest variation (x, y) are shown. The cell-type (a) and pathologist annotations (b, c) are overlaid, with the density of the labels shown across the axes.

**Figure 4:**
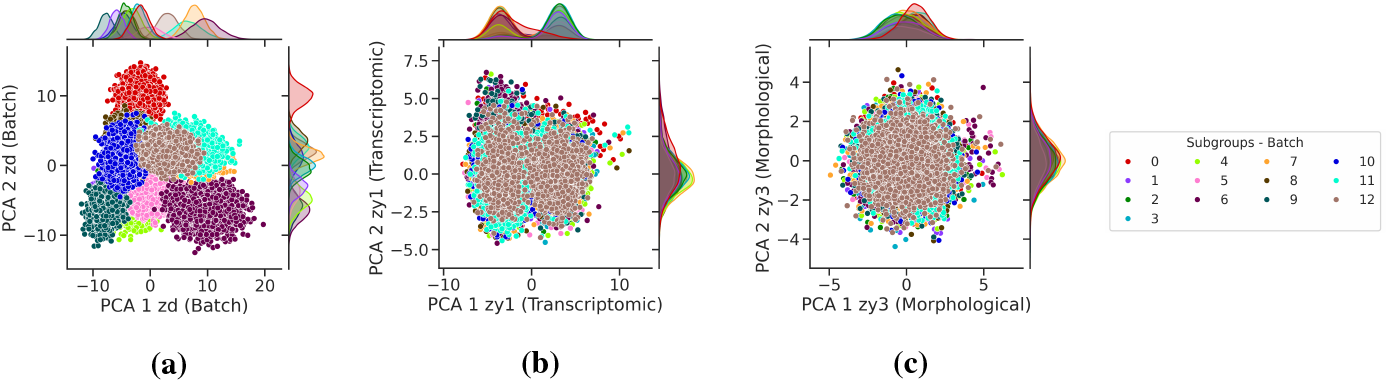
Batch-associated posterior distribution and invariance in other factors. High-dimensional samples from the posterior distributions of the batch-associated (a), transcriptomic-associated (b), and morphology-associated (c) latent spaces, reduced using PCA with the two axes of highest variation presented (x, y). The batch labels, based on the slide number of the samples, are overlaid, with the density of the labels shown across the axes.

Examining the first 2 PCs demonstrates 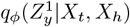 captures variation in the transcriptomic subgroups, specifically separating the fibroblast and pancreatic ductal cell populations (Fig. 3a). The posterior distribution for morphology 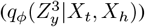 captures distinction between pathologist-annotated tumor/normal areas in the histology data (Fig. 3b). Interestingly, overlaying these labels from the histology data onto the transcriptomics latent space 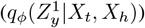 demonstrates the transcriptomics information also distinguishes the tumor and normal pathologist annotations (Fig. 3c), indicating that there is a high degree of mutual information between the pathologist annotations and transcriptomics information in this setting. This type of analysis can be done with any number of factors using SpatialDIVA.

We next investigated how SpatialDIVA removes batch and technical effects (Fig. 1). To our knowledge, SpatialDIVA is the first method that explicitly models batch effects from both the ST expression data and histology image data in a joint manner (Fig. 2a). Batch correction is a challenging task in both the transcriptomics and histology spaces (Kothari et al., 2014; Guo et al., 2023). Visualizing the batch covariate and associated posterior samples (*q*_ϕ_(*Z*_d_ |*X*_t_, *X*_h_)) shows that this latent factor captures the batch variation, as most batches can be distinguished even in the first two PCs of this distribution (Fig. 4a). When we examine the transcriptomic and morphological latent distributions 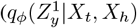 and 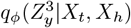 respectively) and overlay the batch/slide labels, we find that this variation has been removed from these factors (Fig. 4b,c), validating that *Z*_d_ effectively captures batch variation.

We compared the batch-correction ability of SpatialDIVA to a conditional VAE (cVAE) model that conditions on batch (slide number), removing all respective across-slide variation (Appendix H). We find that the different posterior distributions of SpatialDIVA minimize batch effects, but do not remove inter-batch variation completely, as biological variation for the relevant posterior distributions 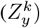 differs between batches (Table 3). As expected, the batch-associated posterior (*Z*_d_) has the lowest batch-correction score, as it captures batch variation (Table 3). The residual variation (*Z*_r_) has the highest score, indicating it is strongly independent of slide context (Table 3). Furthermore, SpatialDIVA preserves biological information in its latent spaces better than the cVAE baseline (Appendix Table 5).

**Table 3:** Batch correction evaluation for the SpatialDIVA model. The average batch silhouette width (*ASW*_batch_) measures the degree of batch mixing where 0 indicates no batch mixing and 1 indicates perfect batch mixing. Results are shown for 5 random seeds for the conditional variational autoencoder (cVAE) model and the different posterior latent distributions of SpatialDIVA. The best results in each dataset are **bolded**.

| Method | Colorectal cancer<br>cohort **batch** | Pancreatic cancer<br>cohort **batch** |
| --- | --- | --- |
|  | **ASW** | **ASW** |
| cVAE | **1.000 ± 0.000** | 0.949 ± 0.114 |
| SpDIVA $Z_y^1$ | 0.729 ± 0.028 | 0.704 ± 0.008 |
| SpDIVA $Z_y^2$ | 0.610 ± 0.031 | 0.393 ± 0.084 |
| SpDIVA $Z_y^3$ | 0.721 ± 0.009 | 0.571 ± 0.018 |
| SpDIVA $Z_r$ | **1.000 ± 0.000** | **1.000 ± 0.000** |
| SpDIVA $Z_d$ | 0.200 ± 0.049 | 0.144 ± 0.020 |

**Table 4:** Disentanglement benchmark results for the pancreatic and colorectal cancer datasets, **standard deviation results from Table 2.** Results are shown from 10 random subsamples of the datasets.

| Metric \ Method | Colorectal cancer cohort |  |  |  | Pancreatic cancer cohort |  |  |  |
| --- | --- | --- | --- | --- | --- | --- | --- | --- |
| | PCA | VAE | β-VAE | SpatialDIVA | PCA | VAE | β-VAE | SpatialDIVA |
| JEMMING | 0.0063 | 0.2730 | 0.0481 | 0.0212 | 0.0098 | 0.2400 | 0.0030 | 0.0254 |
| SAP | 0.0004 | 0.0000 | 0.0000 | 0.0013 | 0.0008 | 0.0000 | 0.0000 | 0.0023 |
| MIG | 0.0005 | 0.0054 | 0.0023 | 0.0020 | 0.0009 | 0.0082 | 0.0025 | 0.0058 |
| MIG-SUP | 0.0012 | 0.0728 | 0.0231 | 0.0019 | 0.0006 | 0.0585 | 0.0181 | 0.0023 |
| Modularity | 0.0051 | 0.0262 | 0.0102 | 0.0077 | 0.0045 | 0.0440 | 0.0311 | 0.0129 |
| DCI-MIG | 0.0044 | 0.0008 | 0.0003 | 0.0166 | 0.0021 | 0.0001 | 0.0001 | 0.0166 |
| Explicitness | 0.0022 | 0.1582 | 0.0971 | 0.0029 | 0.0042 | 0.0581 | 0.0020 | 0.0047 |
| IRS | 0.0014 | 0.0616 | 0.0001 | 0.0016 | 0.0596 | 0.0183 | 0.0000 | 0.0400 |

**Table 5:**
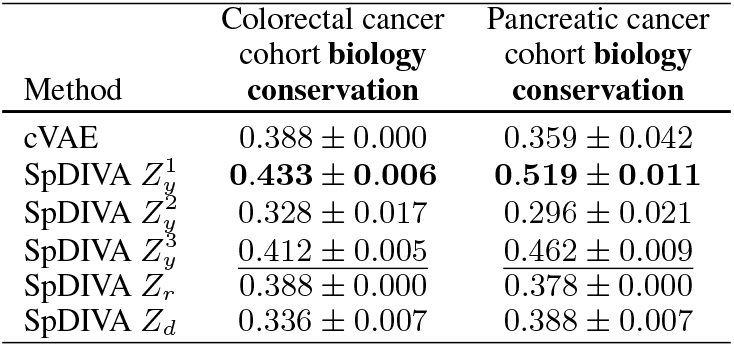
Biological conservation evaluation for the SpatialDIVA model. The average biology conservation score, which is an aggregate of several metrics (Appendix H) is shown. Results are shown for 5 random seeds for the conditional variational autoencoder (cVAE) model and the different posterior latent distributions of SpatialDIVA. The best results in each dataset are **bolded**, and the second-best results are underlined.

### 4.3 Conditional generation of multi-modal data

As SpatialDIVA is a generative model, it is possible to generate new data (*X*_t_, *X*_h_) while intervening on disentangled factors. Specifically, we can sample from the likelihoods by conditioning on specific factors sampled from the posterior, while setting others to a constant value. For instance, if we condition on only transcriptomic context 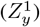:

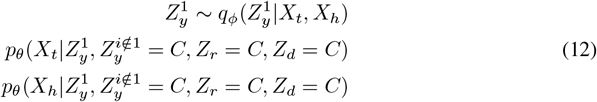

Essentially, Eqn. 12 indicates that the generated samples for *X*_t_ and *X*_h_ will vary according to 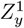, but not the other factors, as they are held constant. This allows us to quantify how one disentangled factor influences variation in the multi-modal data. For example, if we want to evaluate the information that morphology-associated variation 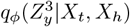 encodes in the ST data (*X*_t_), we can condition on the morphology factor and hold others constant, when generating the transcriptomic likelihood (Appendix Fig. 8).

### 4.4 Conditional generation analysis of PDAC cancer biopsy samples

Using this setup (Sec. 4.3), we sought to understand which gene programs and pathways can be exclusively associated with transcriptomic variation, spatial context, and morphology information, and which are shared, in pancreatic cancer. Specifically, we trained the SpatialDIVA model on all 13 slides of the multi-patient PDAC cohort and used the generated counts for further analysis (Appendix H). After training the model using a negative binomial likelihood for the ST counts (*X*_t_), we sampled the shape (*θ*) and mean (*μ*) parameters for each spot and gene across the PDAC slides, based on each conditional factor: 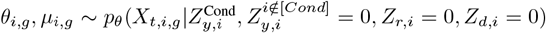. After obtaining these parameters for each spot on a per-gene level, we obtained negative binomial distribution parameterizations which we then sampled counts from: 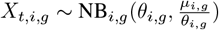. This was done for a random subset of 10000 spots sampled from all PDAC slides.

We first determined whether the differentially expressed genes (DEGs) in the counts generated are specific or shared across factors (Appendix H). Differential expression quantifies which genes exhibit statistically significant variance between clusters of spots, and are indicative of cell-types, cell-states and gene programs that can be captured by transcriptional counts (Wolf et al., 2018) (Appendix H).

The top 500 DEGs in the conditionally generated counts from each factor were almost mutually exclusive (Fig. 5a). This indicates that the gene programs associated with these factors are likely to be mutually exclusive, and is another result that shows the SpatialDIVA model has successfully performed multi-modal disentanglement.

**Figure 5:**
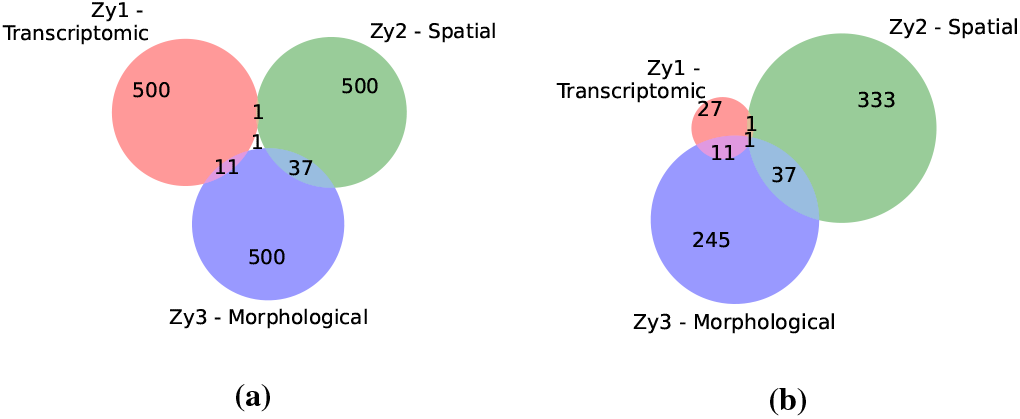
Overlap of DEGs and pathways for conditional generation. For PDAC, conditional generation of the ST counts (*X*_t_) was performed based on the intrinsic transcriptomic 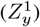, spatial 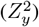 and morphological 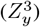 factors of variation. (a) The overlap of the top 500 DEGs for the generated counts conditioned on each factor. (b) Overlap of the enriched pathways in the top 500 DEGs for each factor.

We analyzed more concretely the higher-level gene programs encoded by the top DEGs based on pathway enrichment analysis. Pathway analysis assesses statistical overrepresentation for a set of genes in biological pathways curated by experts (Kolberg et al., 2023) (Appendix H). As expected based on the DEG analysis, the overlap of the enriched pathways based on each conditioned factor was minimal (Fig. 5b). Further examination of the pathways based on keyword overlap (Appendix H) reveals that intrinsic transcriptomic variation 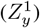 encodes general pancreatic functions (Fig. 6a).

**Figure 6:**
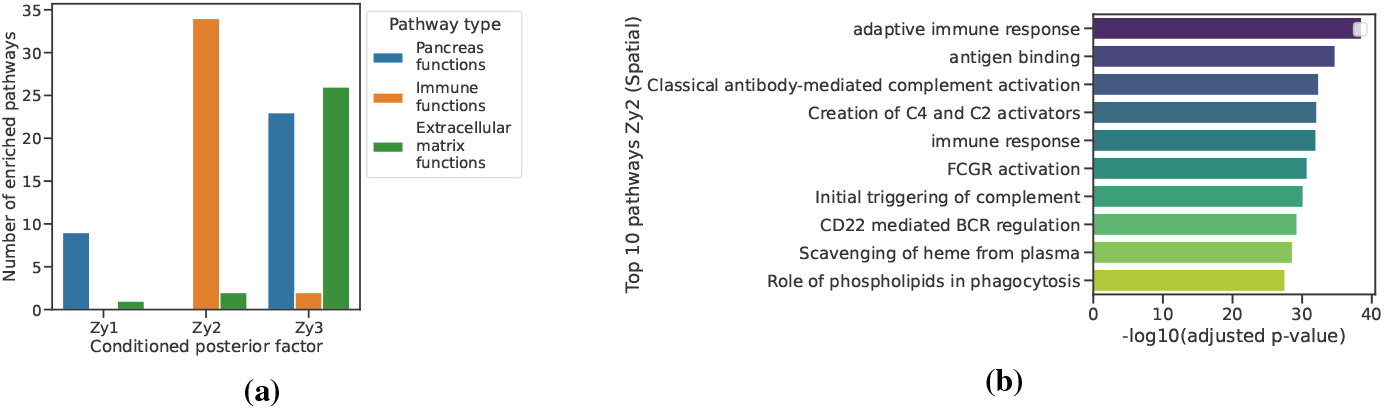
Pathway enrichment of top 500 DEGs from conditional generation. (a) The number of enriched pathways specific to a functional group, from the generated counts conditioned on each factor. (b) The top 10 pathways with the highest enrichment based on corrected *p*-value for the 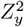 (spatial) conditioned generated transcriptomic counts.

Spatial variation 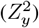 was found to capture the majority of the immunological signal (Fig. 6a,b). This result is important as PDAC characterization and progression is significantly influenced by immune cells in the tumor microenvironment (Karamitopoulou, 2019). The morphological variation 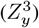 was enriched for pancreas-specific and extracellular matrix-specific functions (Fig. 6a).

PDAC has been subtyped extensively, and the general consensus is that there are two transcriptionally distinct subtypes - classical and basal presentations (Moffitt et al., 2015). These subtypes have unique histological, genomic and transcriptional presentations, and basal-like PDAC is associated with worse prognosis overall (Moffitt et al., 2015; Singh et al., 2024). We sought to determine which subspaces in the PDAC-trained SpatialDIVA model are associated with classical and basal transcriptional marker genes. To do so, we took a list of marker genes for the classical and basal presentations (Appendix H), and determined the overlap of the top 500 DEGs from each conditionally generated subspace (Fig. 7). Even though the conditional analysis considers all spots (and hence all cells) across PDAC biopsies, and not exclusively tumor cells, we observed a strong signal for both the basal and classical markers in the DEGs of the intrinsic transcriptomic, spatial, and morphology-associated subspaces (Fig. 7). For the basal subtype, variation associated with spatial context had the highest signal, while the classical subtype was most prevalent in the morphological subspace. These results indicate that variation associated with the distinct PDAC subtypes cannot be predicted by the same generative factors in the multi-modal ST and histology data.

**Figure 7:**
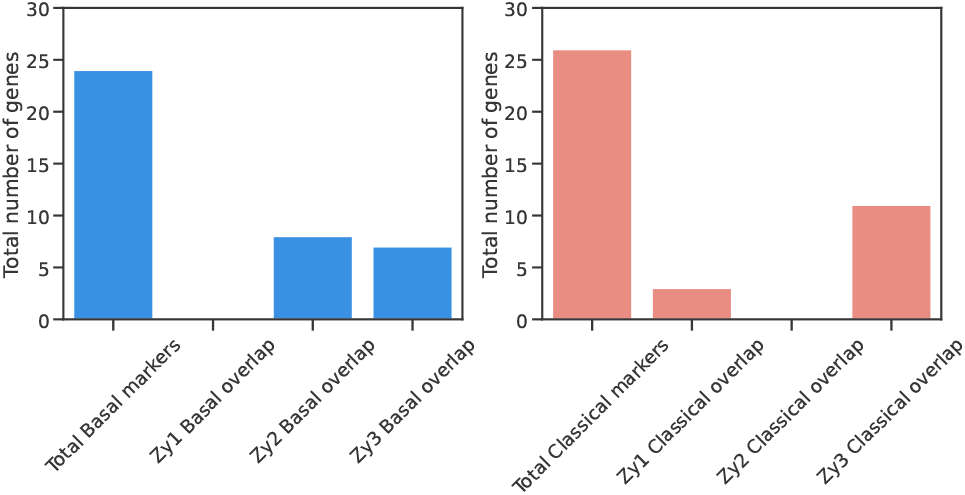
PDAC subtype marker gene enrichment of top 500 DEGs from conditional generation. In each of the plots, the total number of literature-derived marker genes for the basal (left) and classical (right) subtypes is indicated. The overlap of the top 500 DEGs derived from the conditionally generated transcriptional counts (*X*_t_) for the intrinsic transcriptomic 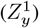, spatial 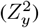, and morphological 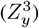 subspaces is shown for each of the subtypes, as an overlapping number of genes with their respective marker gene sets (y-axis).

**Figure 8:**
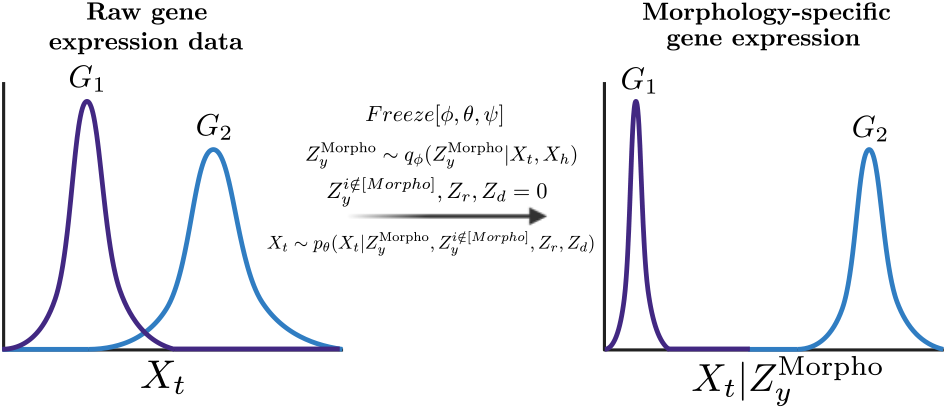
Conditional generation of transcriptomic data. The trained model can be frozen and posterior samples for the morphology-specific latent can be used to generate morphology-specific ST gene expression values, as the other factors are held constant (0) during generation.

This analysis offers a vignette showing the capabilities of SpatialDIVA in determining the contribution of disentangled factors to specific information in the observed data. Depending on the clinical importance of certain gene programs, specific assays and analyses can be prioritized, such as assays for spatial variation, which was found to capture most of the immunological and basal-like subtype signal in the PDAC data.

## 5 Related work

### ST and H&E models

Existing task-specific models for ST data include infering cell-cell communication in a given tissue context (Fischer et al., 2023), general representation learning (Xu et al., 2024), deconvolving cell-types in ST spots (Ma & Zhou, 2022), and aligning ST data with single-cell RNA sequencing data (Biancalani et al., 2021).

Advances in foundation models have naturally led to an increasing interest in their uses for biological data (Consens et al., 2023). Large pretrained models for histopathology data have been developed since the rise of self-supervised pretraining in computer vision models (Ciga et al., 2022; Chen et al., 2024; Xu et al., 2024). In our work, we use the UNI model (Chen et al., 2024) to obtain informative representations of the histology images (*X*_h_). ST foundation models have also been introduced (Schaar et al., 2024; Cao & Yuan, 2024; Wang et al., 2025), and can be used to generate embeddings of the ST counts (*X*_t_) for use in the SpatialDIVA model.

Several models have been developed that aim to learn joint representations of the ST and histology data, for various downstream tasks (Hu et al., 2021; Xu et al., 2022; Pham et al., 2023; Zhao et al., 2024). However, these models do not learn disentangled representations of factors of variation for both modalities, as we do in this work.

### Histology to gene expression prediction models

Many models exploit paired ST and histology to predict gene expression for a given region of an H&E slide (He et al., 2020; Pang et al., 2021; Zeng et al., 2022; Xie et al., 2023). Although this approach can impute gene expression from morphology alone, it is bounded by the maximum mutual information between the two modalities and does not return any relevant information on the specific type of information in each, as well as their associated generative factors.

### Disentangled representation learning

There have been several advances in unsupervised disentanglement (Higgins et al., 2016; Chen et al., 2018). However, Locatello et al. (2018) demonstrated empirically and theoretically that without inductive biases, there are no guarantees for learning disentangled representations. Consequently, multiple methods adopted strong and weak supervision, as well as semi-supervised approaches (Ilse et al., 2019; Joy et al., 2020; Locatello et al., 2020; Brehmer et al., 2022).

The two approaches that form the basis for our work are the **domain invariant variational autoen-coder (DIVA)** (Ilse et al., 2019) and the **characteristic capturing VAE (ccVAE)** (Joy et al., 2020) models. Although derivatives of these works have been applied to biological data (Pradier et al., 2023), the models were not designed to handle multi-modal data (including ST and histology) and continuous label distributions, as is the case for our spatial supervision covariate. As such, we explicitly build upon DIVA and ccVAE for disentanglement of spatial transcriptomics and histopathology imaging.

## 6 Conclusion

Here, we framed the challenge of determining the generative factors of ST and histopathology, and their overlap, as multi-modal disentanglement. We introduced *SpatialDIVA*, the first technique to perform multi-modal disentanglement in this setting, which leads to an interpretable and flexible set of posterior distributions that are able to generate novel biological insights through multi-modal conditional generation. The problem formulation and evaluation framework we developed is an important resource for the community in creating models that learn factors relevant to each modality.

## Acknowledgments

The authors thank Michael Geuenich and Chengxin Yu for providing curated lists of PDAC marker genes. HM thanks Julia Greissl, Paidamoyo (Ash) Chapfuwa, Ted Meeds, Melanie F. Pradier, and Niranjani Prasad from Microsoft Research for insightful discussions on DIVA and related models.

## A SpatialDIVA - Extended Background and Problem Formulation

### A.1 Biologically informative priors

Through outlining the flow of information within and across cells, we can determine some of the important factors that generate the ST and histology data distributions.

#### Intrinsic transcriptomic variation

Variation in the RNA expression of different genes caused by cell intrinsic genetic or epigenetic factors, is an important driving factor for both the ST and histology data. This affects both ST expression variation as well as morphological variation, driven by changes in expression of morphology-associated genes (Kiger et al., 2003).

#### Morphological variation

Morphological variation on a per-cell level clearly influences the morphological variation on a per-spot level. However, changes in protein expression can also influence transcriptomic variation (Vogel & Sheetz, 2006; Dupont et al., 2011) - there is not a simple linear flow of information from DNA to RNA to protein expression. Therefore, this factor of variation can affect both ST and histology readouts.

#### Spatial variation

Aside from intrinsic cellular variation at the per-spot level, variation in both transcriptomic and morphological distributions can be driven by spatial context (Bressan et al., 2023). The most concrete example of this is cell to cell communication across spatial domains. This type of communication can influence transcriptomic variation, which in turn can affect both the ST and histology contexts.

#### Technical/batch variation

Biological factors can influence changes in spot-level representations of the data, however, technical variation can also drive these changes. Batch effects are variations caused by assay differences, experimental differences, or even ambient conditions (Leek et al., 2010). This variation is often conflated with biological variation, unless specifically accounted for (Leek et al., 2010).

#### Residual variation

Any variation in the ST and histology data distributions that is not accounted for by the aforementioned factors is considered residual variation. If batch/technical effects have been accounted for, residual variation should capture biological effects not captured by the prior information injected through the other factors.

## B SpatialDIVA - extended results

### B.1 Results Section 4.1

### B.2 Results Section 4.2

### B.3 Results Section 4.3

## C Datasets and dataset formatting

The SpatialDIVA model is dependent on having spot-level alignment of ST data and patches from the histology image. Given this constraint, we used data that was either already processed by the HEST-1k pipeline (Jaume et al., 2024), or used their preprocessing scripts to process other paired ST/histology data.

### C.1 ST data preparation

#### C.1.1 Colorectal cancer data

The spatial transcriptomics data for the colorectal cancer cohort (Valdeolivas et al., 2024) was obtained from the HEST-1k dataset of jointly measured ST and histology data using the Visium platform (Jaume et al., 2024). The HEST-1k uses a specific pipeline for segmentation of tissue and alignment of ST and histology data (Jaume et al., 2024). Given the data as it was procecessed by HEST-1k, we aimed to match the annotations by the authors in the original paper for the ST spot-level cell-type deconvolution and expert pathologist annotations (Valdeolivas et al., 2024), to the data as processed by HEST-1k. We used the barcode information from each ST sample to achieve this, and subset the data from HEST-1k to spots that have both pathologist and ST cell-type annotations from the supplementary information from Valdeolivas et al. (2024). This resulted in 14 tissue sections, corresponding to 14 samples for paired ST and histology measurements.

#### C.1.2 Pancreatic cancer data

The pancreatic cancer data (Cui Zhou et al., 2022) was reprocessed and reannotated with updated single-cell references (Peng et al., 2019; Yu et al., 2024).

The dataset consists of 15 sections from 10 patients available on the Human Tumour Atlas Network (HTAN) under the atlas code WUSTL. To estimate the number of cells per spot, we performed nuclear segmentation on the full resolution histology image. Briefly, image crops of each Visium spot were obtained and nuclear segmentation was performed using squidpy (Palla et al., 2022) and stardist (Schmidt et al., 2018) to obtain cell counts per spot. An average cell count per spot was derived for each Visium slide after segmentation of all spots. To infer cell-type proportions for each spot, we utilized cell2location (Kleshchevnikov et al., 2022) and cell-type labels from publicly available single cell adenocarcinoma datasets from Peng et al. (Peng et al., 2019) and Cheng et al (Yu et al., 2024). Briefly, the single cell reference was trained in cell2location using the following parameters for gene selection: (cell count cutoff: 5, cell percentage cutoff2: 0.03, nonz mean cutoff: 1.12, non-mitochondrial genes) and training (num samples: 1000, batch size: 2500, num epochs: 250, batch key: sample id). To predict cell-type proportions for each Visium sample, we utilized the average cells per spot from segmentation for each slide and the following parameters (detection alpha: 20, max epochs: 30000). Lastly, the 5% quantile of the posterior distribution was utilized as the inferred cell-type abundance per spot and projected onto the histology image for visualization. We used 13 of the 15 sections due to annotation limitations for the histology data (Sec. C.2).

For both the colorectal and pancreatic cancer data ST-labels, which are a proportion of cell-types per spot, we selected the cell-type that had the highest proportion in each spot to use as a categorical label when training and evaluating the SpatialDIVA model.

End-users of the SpatialDIVA method can use their own datasets for ST data, provided there are labels for both the cell-type (ST-derived) and pathologist annotations. In cases where these labels have not yet been derived, unsupervised clustering can be used within each modality to obtain labels.

### C.1 Histology data preparation

Alignment of ST data at the level of spots and the histology data which is split up into patches, is necessary for training the SpatialDIVA model. The Visium data from HEST-1k (Jaume et al., 2024) was processed using an end-to-end pipeline that the authors developed for Visium datasets that performs automatic tissue segmentation, alignment, and resolution detection. The pipeline results in processed ST data with a measurement of gene expression per spot on the ST slide, and histology patches that are centered around each ST spot. The pipeline does this by creating 224×224 px patches at 20X magnification around each spot. This results in patches of histology images that approximately correspond to each of the ST spots in the slide.

From here, we extract image features for each spot-aligned patch using the UNI foundation model (Chen et al., 2024). The UNI model is loaded via the timm library (Wightman, 2019) and the Huggingface Hub (Wolf et al., 2019). The pretrained model is loaded via the following parameters: {pretrained = True, init values = 1e-5, dynamic img size = True}. Specifically, we used the ViT-L/16 model from the first version of UNI (Chen et al., 2024). Using UNI in inference mode with no further fine-tuning, we obtained 1024 dimensional embeddings for each spot-level patch for both the colorectal cancer and pancreatic cancer datasets.

#### C.2.1 Colorectal cancer data

The colorectal cancer data (Valdeolivas et al., 2024) had already undergone the respective processing using the HEST-1k pipeline to obtain spot-level patch representations of the histology image. As such, there was no further processing necessary and we used these patches and the corresponding ST spots (Sec. C.1) with ST cell-type labels and pathologist annotations from the original study (Valdeolivas et al., 2024) for further analysis. The pathologist annotations were done at the spot-level, so aligning these with the spot-level patches used the same code as aligning cell-type labels.

#### C.2.2 Pancreatic cancer data

The pancreatic cancer data (Cui Zhou et al., 2022) was not present in the HEST-1k dataset. As such, we used the HEST-1k pipeline to perform tissue segmentation, upscaling, and patching at the spot-level for this data (Jaume et al., 2024). Specifically, we used the VisiumReader() function from HEST-1k, which loads the histopathology image, feature matrix for ST, and the spatial coordinates. Then we used a built-in function for OTSU segmentation, followed by patching at a size of 224 px. This resulted in patches corresponding to ST spots for the segmented tissue from HEST-1k. The ST data that we had from (Cui Zhou et al., 2022) was inner-joined with the spots that were segmented through the HEST-1k pipeline.

For the samples of the pancreatic cancer data (Cui Zhou et al., 2022), of the 15 samples, 13 of them were annotated by a clinical pathologist in our team, for tumor versus normal regions of the slide. As such, we only used the 13 pathologist-annotated samples from the original study for subsequent analysis. The pathologist annotations for the 13 samples were transferred onto the ST spots through the Shapely library (Gillies et al., 2007).

After processing both datasets, the resulting representations had spot-aligned UNI features (1024 dimensional), pathologist annotations at the spot-level, deconvolved cell-type annotations at the spot-level, as well as expression across all mapped genes, spatial coordinates on each slide for each spot, and other metadata that was used (such as batch/slide number).

## D ST and histology data preprocessing

### D.1 ST data preprocessing

After data preparation, the raw counts from the ST data can be processed further depending on the experiments.

In general, for the ST-derived labels, which comprised of deconvolved cell-type proportions for each spot, we took the cell-type with the highest proportion (representation) for each spot as a soft categorical label. This label was used for model training and evaluation across all results sections.

For evaluation of the SpatialDIVA model versus baselines in quantification of disentanglement (Results Sec. 4.1, Table 2) and the quantitative evaluation of batch-correction effects (cVAE and SpatialDIVA, Results Sec. 4.2, Table 3), the SpatialDIVA model and baselines were trained after the following preprocessing procedures to the ST counts:

- Count normalization per spot to a fixed value (10000)
- Log1p, equivalent to *ln*(*x* + 1), transformation of the counts
- Highly-variable gene selection using the ‘seurat’ method

Count normalization and Log1p transformations were done for each sample/slide individually, and samples for the complete datasets (pancreatic and colorectal cancer) were concatenated to perform a joint highly-variable gene selection. Joint highly-variable gene selection ensures that cell-types and states across all slides are best captured. These transformations were done using the Scanpy library (Wolf et al., 2018).

For the qualitative disentanglement results in Figs. 3 and 4, the SpatialDIVA model was trained after count and Log1p normalization, but no highly-variable gene selection was done and all genes were used in training the model and evaluating the empirical posterior distributions.

### D.2 Histology data preprocessing

The extracted UNI (Chen et al., 2024) features for the spot-level patches of histopathology image data (Sec. C.2) were standardized at the feature-level (1024 dimensions = 1024 features) for all experiments. Standardization of these features was done for each slide (sample) individually. The reasoning for this was to ensure that batch-effects are not introduced by standardization of the UNI features across samples/slides (Lin & Lu, 2022).

### D.3 Preprocessing and experimental setup

A potential challenge in the across-slide selection of highly-variable genes and within-slide standardization of UNI features, is when evaluating disentanglement quantitatively (Sec. 4.1). However, we consider all samples of one cohort to be part of a single observed data distribution, and effectively evaluate samples that are i.i.d. in the disentanglement benchmark. As such, factors such as generalization and data leakage are not considered, because this is a statistical learning problem. This is in line with previous work on evaluating disentanglement (Locatello et al., 2018). The approach of training and evaluating on different subsets of a known data distribution can be interpreted as bootstrapping our estimates of disentanglement performance (Sec. 4.1).

## E SpatialDIVA model details

### E.1 SpatialDIVA distributions

The SpatialDIVA model architecture is that of a neural network with separate encoders for each posterior covariate (*q*_ϕ_(*Z*_i_)) that is considered in the model, linear decoders that use samples from the posterior distribution to reconstruct the likelihoods for the ST (*X*_t_) and histology (*X*_h_) data distributions (*p*_θ_(*X*_i_)), encoders for the prior distributions (*p*_θ_(*Z*_i_)), and classification heads for the label distributions (*q*_ψ_(*L*^i^)). Table 6 summarizes the distributions used in the model, as well as the data required for each.

**Table 6:**
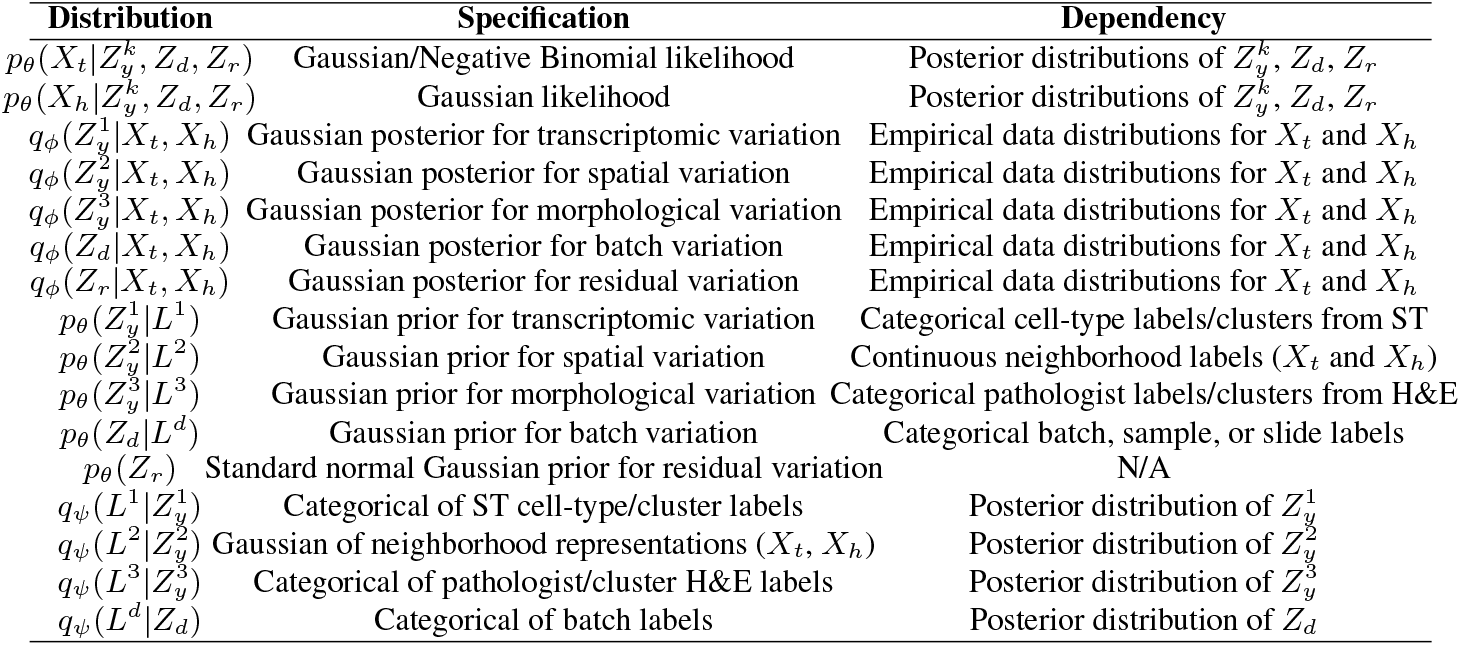
Distributions used in the SpatialDIVA model and details.

### E.2 SpatialDIVA architecture

For each of the given distributions outlined in the previous section, Tables 7, 8, 9, 10, and 11 summarize the SpatialDIVA architecture choices for the different experiments in the paper. Exceptions based on experiments are noted after the tables.

**Table 7:** Architecture for data likelihood distribution for ST – 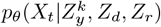.

| Module number | Component |
| --- | --- |
| 1 | nn.Linear(100, 64) |
| 2 | mu = Softplus(nn.Linear(64, 36601)) |
| 3 | logvar = nn.Linear(64, 36601) |

**Table 8:** Architecture for data likelihood distribution for histology - 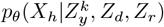.

| Module number | Component |
| --- | --- |
| 1 | Linear(100, 64) |
| 2 | mu = Linear(64, 1024) |
| 3 | logvar = Linear(64, 1024) |

**Table 9:** Architecture for posterior distributions - *q*_ϕ_(*Z*_i_|*X*_t_, *X*_h_)

| Module number | Component |
| --- | --- |
| 1 | Linear(37625, 64) |
| 2 | ReLU() |
| 3 | mu_zy = Linear(64, 20) |
| 4 | logvar_zy = Linear(64, 20) |

**Table 10:** Architecture for prior distributions - *p*_θ_(*Z*_i_ | *L*^i^). **Variable** here indicates the variable input length for the prior labels, which depends on the posterior/label combination and dataset.

| Module number | Component |
| --- | --- |
| 1 | Linear(**Variable**, 64) |
| 2 | ReLU() |
| 3 | mu_zy = Linear(64, 20) |
| 4 | logvar_zy = Linear(64, 20) |

**Table 11:**
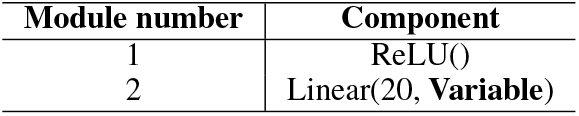
Architecture for label distributions - *q*_ψ_(*L*^i^ | *Z*_i_). **Variable** here indicates the variable output length for the prior labels, which depends on the posterior/label combination and dataset.

In terms of **exceptions** to the architectures outlined in Tables 7, 8, 9, 10, and 11:

- For the experiments benchmarking disentanglement (Table 2) and batch-correction (Table 3), the input size for ST counts (*X*_t_) was 2500 (instead of 36601), as highly-variable gene selection was done for these experiments (changes Table 7 output size, Table 9 input size)
- For the latent space covariance analysis, the dimensionality of output for the posterior of batch variability (*q*_ϕ_(*Z*_d_ | *X*_t_, *X*_h_)) was changed from 20 to 5 (Table 9). Also for this analysis, two hidden layers were used for the ST and histology data distributions (*p*_θ_(*X*_t_ |..), *p*_θ_(*X*_h_ |..)) of sizes 256 followed by 128 (Tables 7 and 8). Lastly, an extra hidden layer was used for the encoders of the posterior distributions (*q*_ϕ_(*Z*_i_ | *X*_t_, *X*_h_)), of size 32 (two sequential hidden layers, size 64 and 32 with ReLU() non-linearities after each) (Table 9)
- For the conditional generation experiments (Results Sec. 4.3 and 4.4), the likelihood distribution for ST (Table 7) outputs *θ* and *μ* to parametrize a Negative Binomial distribution, which are both constrained to be positive via Softplus, instead of *μ* and logvar for a Gaussian distribution. Further, the dimensionality of output for the posterior of batch variability (*q*_ϕ_(*Z*_d_ | *X*_t_, *X*_h_)) was changed from 20 to 5 (Table 9). Two hidden layers were used for the ST and histology data distributions (*p*_θ_(*X*_t_ |..), *p*_θ_(*X*_h_ |..)) of sizes 256 followed by 128 (Tables 7 and 8). Lastly, an extra hidden layer was used for the encoders of the posterior distributions (*q*_ϕ_(*Z*_i_ | *X*_t_, *X*_h_)), of size 32 (two sequential hidden layers, size 64 and 32 with ReLU() non-linearities after each) (Table 9).

## F SpatialDIVA objective derivation

For the SpatialDIVA model (Fig. 2), we want to maximize the following marginal likelihood of the ST counts (*X*_t_), histology features (*X*_h_) and observed labels (*L*):\

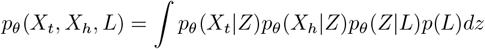

However, this marginalization over all possibilities of *Z* is intractable. We can instead learn a lower bound on the log-likelihood:

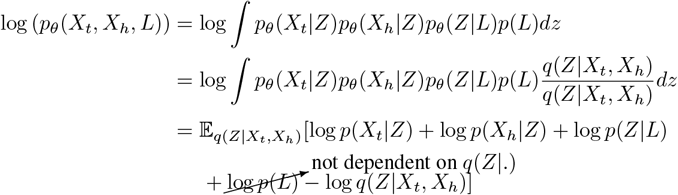

Using Jensen’s inequality, which indicates that log(E(..)) ≥ E[log(..)]:

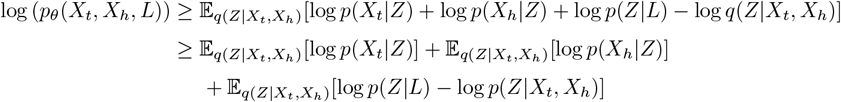

We have two types of label distributions - *L*^k^ for *k* biologically informative prior labels, *L*^d^ for the batch/sample. These correspond to 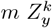 posterior distributions, a *Z*_d_ posterior. There is also a posterior for residual variation in *Z*_r_. We can break down this bound further based on these distinct factors, without explicitly summing over the *k* posteriors and labels for biologically informative labels. The posterior for residual variation (*Z*_r_) is not constrained by a label distribution.

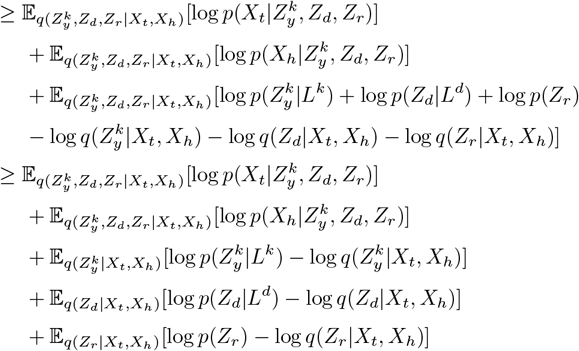

Through the definition of the KL-divergence:

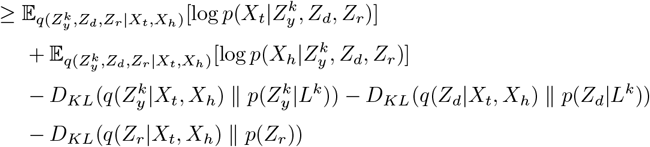

The first two expectations correspond to the likelihoods of the ST (*X*_t_) and histology (*X*_h_) data. The last three KL divergence terms penalize the learned posterior distributions based on the learned prior distributions. The exception is *q*(*Z*_r_ *X*_t_, *X*_h_), which is penalized based on a standard normal Gaussian prior *p*(*Z*_r_). We can decompose the KL divergences for all of the *Z*_y_ terms we consider, including intrinsic transcriptomic variation 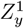, spatial variation 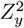, and morphological variation 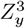:

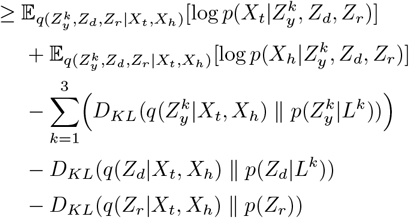

Similar to ccVAE (Joy et al., 2020) and DIVA (Ilse et al., 2019), we incorporate classification losses for the posterior samples based on the labeled data (Appendix 6):

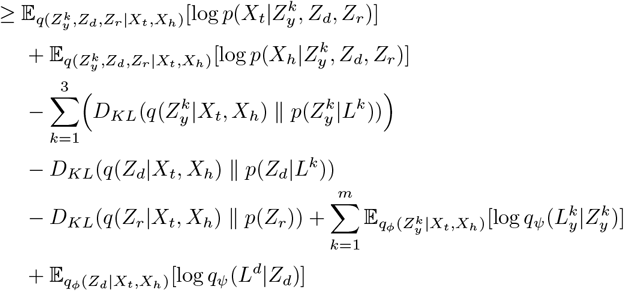

This leads to the following objective, with *β*’s corresponding to hyperparameters. We minimize the loss, and hence the signs flip:

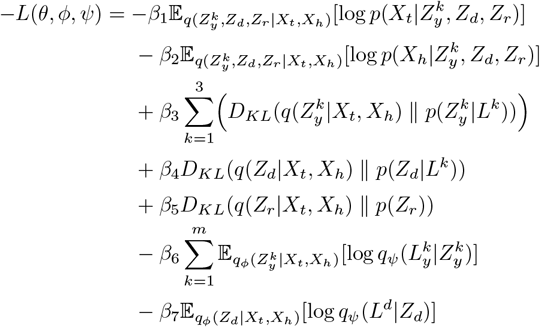

The parameterizations of *θ, ϕ*, and *ψ* for the different distributions are outlined in Appendix E.

In general, across experiments, we use hyperparameters where all *β*’s are set to 1. Exceptions are explicitly indicated in Appendix H.

## G Biological factors of variation in ST and histology

For the key biological factors of variation that we considered in our analysis that are initially described in Appendix A.1, we provide more detail on the rationale behind their selection, framing in the problem, and preprocessing below.

### G.1 Intrinsic transcriptomic variation – 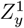

Intrinsic variation of gene expression takes into account the biological processes that are not strongly influenced by cellular organization and cell-cell communication. Within a tissue context there is a large-scale orchestration of cell-cell communication that occurs via signalling, both short and long-range (Alberts et al., 2022). Even longer range signals can arrive from entirely different organs and parts of the body that influence the transcriptomic state of individual cells, for functional effects such as differentiation (Alberts et al., 2022). Variation in gene expression that is intrinsic, therefore, should capture effects and functional changes to gene expression that are dictated by the internal state of a cell. An example of this could be immune evasion functions governed by gene expression changes that are activated in cancer cells due to mutations (Hanahan, 2022).

We consider intrinsic transcriptomic variation to be an important factor, as this variation will affect both the transcriptomic counts (*X*_t_) in a trivial manner, and the histology features (*X*_h_) due to the relationship between transcription and protein expression (more details provided in the Morphology section). A potential challenge is determining how to isolate intrinsic versus extrinsic transcriptomic variation, where extrinsic variation is due to factors such as cell-cell communication. From a supervision perspective, the labels we utilize, whether they are clusters or expert-annotated cell-types derived from the ST data, will also likely be influenced by extrinsic variation as this will affect the ST counts (*X*_t_) as well. In SpatialDIVA, we aim to remove the extrinsic variation captured in the posterior distribution for 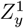 by introducing a spatial variation covariate 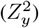. Although we constrain intrinsic transcriptomic variation on labels that may contain information from both intrinsic and extrinsic sources, by encouraging disentanglement through the 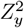 posterior, we aim to remove as much extrinsic variation from the posterior distribution of 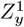 as possible.

In our analysis, we use expert-annotated cell-type labels derived from the ST counts, for both the colorectal and pancreatic cancer datasets (*L*^1^). These labels are derived using a single-cell reference, as the Visium ST protocol does not yield single-cells per spot but a mixture of cells (Ståhl et al., 2016). The process is referred to as cell-type deconvolution, and returns a proportion of cell-types estimated to be in each spot of the ST slide (Ma & Zhou, 2022). As we use a categorical distribution of labels to constrain our posterior for intrinsic transcriptomic variation (Appendix 6), we take the cell-type that has the *highest proportion in each spot as a label*, and we consider the entire set of cell-type labels derived in this manner as a categorical distribution. This was done for both the colorectal cancer data and re-annotated pancreatic cancer datasets.

### G.3 Spatial variation – 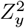

Spatial variation considers the influence of extracellular signalling as well as more direct means of cell-cell communication, such as at cellular junctions (Alberts et al., 2022). Spatial context is relevant across many biological scenarios, including development and cancer. Within development, cellular signalling gradients which are organized spatially, dictate how certain cells differentiate and what cells and tissues they will give rise to (Barresi & Gilbert, 2023). In cancer, as we highlight in the case of PDAC, spatial context is important for characterization of a tumor as well as potential diagnostic and therapuetic avenues (Karamitopoulou, 2019; Cui Zhou et al., 2022). We consider both intrinsic transcriptomic and morphological variation as separate covariates, and therefore, the spatial variation posterior 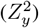 should aim to capture spatial variation that will affect both the ST (*X*_t_) and histology profiles (*X*_h_). Therefore, for our spatial context covariate (Eqns. 10, 11), we consider the concatenation of reduced features for both the transcriptomic (*X*_t_) and histology data (*X*_h_).

Essentially, through our neighborhood decomposition, we aim to represent a label distribution (*L*^2^) that captures the *context* of all the cells in each spot *i*, by having a representation that is predictive of the *X*_t_ and *X*_h_ features of the spatial neighbors of *i*. These neighbors are defined by locality, and we are using the spatial coordinates available in the data (*P*_x,y_) to create this predictive representation. If the prior and posterior distributions for 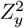 retain this context, and if disentanglement in the model is working appropriately, the spatial context should be captured by 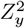, while being minimized in all of the other posterior distributions.

### G.3 Morphological variation – 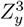

The last factor of variation that we considered was morphological variation. Cellular morphology is dictated primarily by protein expression, cellular shape, and organization of subcellular organelles like the nucleus (Alberts et al., 2022). Morphological features have a long history of being utilized to differentiate the state of cells, such as the use of H&E staining in histopathology to differentiate cancer cells from normal cells (He et al., 2012). Morphological features are distinguished through the histology aspect of the multi-modal data we consider (*X*_h_), and therefore have a direct effect on this modality, much like intrinsic transcriptomic variation has a direct effect on the transcriptomic counts per spot (*X*_t_). However, as described in Appendix A.1, both morphological and intrinsic transcriptomic variation can affect both the transcriptomic (*X*_t_) and histology (*X*_h_) readouts.

Further, we have observed a direct case where transcriptomic variation is correlated with morphological features, in that intrinsic transcriptomic variation 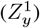 was found to be predictive of a histopathology derived label (tumor/normal) in PDAC (Fig. 3).

For labels that constrain the morphology posterior and prior distributions (*L*^3^), we used pathologist annotations of the histopathology slides based on regions. For the colorectal cancer data (Valdeolivas et al., 2024), this comprised of annotations for distinctive tumor, stromal, and normal regions of cells stained by H&E. For the pancreatic cancer data (Cui Zhou et al., 2022), the slides were annotated by a pathologist on our team, delineating tumor and normal epithelial regions. As such, we had a categorical distribution of labels for the colorectal cancer data, and a binary distribution of labels for the pancreatic cancer data. These labels were derived exclusively through the histology features, and they should thus be significantly correlated with the morphological variation we aimed to capture in this covariate 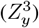.

## H Experiment details and configurations

This appendix section provides details on the experiment settings, as well as baselines that were used and their setup. The different experiments and their details are contained in their own sections.

### H.1 Results Sec. 4.1

#### H.1.1 Quantitative disentanglement benchmark

For the quantitative evaluation of disentanglement, we used a modified version of SpatialDIVA with highly-variable genes (Sec. E.2). For both the colorectal and pancreatic cancer datasets, preprocessing was done in a uniform manner for all baselines and the SpatialDIVA method:

- ST count (*X*_t_) normalization to 10000 counts per spot
- Log1p transformation per spot-gene pair across all spots and genes
- Highly-variable gene selection using all slides in a cohort (2500 genes)
- Standardization of UNI features (1024) for the histology (*X*_h_) data, per slide

For both the colorectal cancer and pancreatic cancer datasets, we sampled 90% of the spots from the combined slides and held-out 10% for evaluation of disentanglement. These splits were done randomly using the numpy (Harris et al., 2020) library, and 10 random seeds.

The **baselines** for disentanglement benchmarking were set up as follows:

**PCA**: For principal component analysis, the transcriptomic counts (*X*_t_) and standardized UNI histology features (*X*_h_) were concatenated and the transcriptomic counts were also standardized. A PCA reduction (Pedregosa et al., 2011) was done on the concatenated representation, and the first 20 dimensions were used as embeddings for disentanglement quantification.

**VAE**: For a base variational autoencoder, we considered a VAE (Kingma & Welling, 2013) with a 20 dimensional latent posterior (*q*_ϕ_(*Z* | *X*_t_, *X*_h_)), with a 64 dimensional hidden layer for the encoder and decoder. ReLU activations were done after the hidden layers, but not before the likelihood and posterior mean and logvar output steps. Similar to SpatialDIVA, a Gaussian posterior and likelihood were used.

*β***-VAE**: The configuration for the *β*-VAE and VAE were the exact same, the only difference was that the weight on the KL divergence term for the posterior distribution was increased to 100, as outlined in Higgins et al. (Higgins et al., 2016):

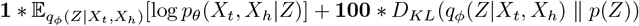

**SpatialDIVA** used the configuration outlined previously, with a Gaussian likelihood for the log-normalized transcriptomic counts (*X*_t_) as well as the standardized UNI histology features (*X*_h_). The trained SpatialDIVA model is used on the test data (frozen model) to extract the mean parameters (*μ*_i_) from the following posterior distributions: 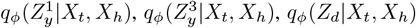. The means from these distributions were then concatenated to obtain test embeddings.

The **PCA** baseline was trained directly on the training set for each iteration using a singular value decomposition (SVD) solver (Pedregosa et al., 2011). The test data is then projected onto the principal components (n=20) calculated via the training data. This 20 dimensional embedding is used for further testing.

The **VAE** and *β***-VAE** baselines were trained on each training subset and the test embeddings from the latent spaces (*q*_ϕ_(*Z*_d_ | *X*_t_, *X*_h_)) were extracted after the models were frozen. The 20 dimensional means *μ*_i_ of the latent embeddings were used for further testing.

The **VAE**, *β***-VAE, and SpatialDIVA** models were trained using the Adam optimizer (Kingma & Ba, 2014) at a learning rate of 0.001 for 100 epochs and batch size of 64.

The extracted embeddings for **SpatialDIVA** and each of the **baseline** methods were used to compute the disentanglement metrics (Appendix I). The factors used in the metrics comprised of the **cell-type labels (from the ST data), pathologist annotations from the histology, and batch labels**. Essentially, these metrics measure how well the latent spaces of the baselines, as well as the SpatialDIVA method, capture variance in ST-labelled cell-types, pathologist-annotated histology data, and batch/technical effects. These factors were one-hot encoded and were used with the embeddings extracted from each method to quantify the disentanglement scores across metrics (Appendix I).

The assessment of disentanglement for this setup was done for both the pancreatic and colorectal cancer datasets (Table 2).

Using the metric values, the methods were *ranked* based on performance, where a 1 indicated the best rank in a dataset for a given metric and 4 indicated the worst rank. Ranks were added up across metrics for the methods and the method with the lowest aggregate score was the best and was ranked first overall, and the other methods followed and were ranked using the same aggregation procedure.

### H.2 Results Sec. 4.2

#### H.2.1 Disentangled latent space and covariance analysis

To assess the latent spaces of the posterior distributions, the SpatialDIVA model was trained on all of the genes quantified in the ST (*X*_t_) data for the pancreatic cancer cohort, with all slides used for training the model. In terms of architecture, the architecture indicated in Sec. E was used with the indicated changes (Sec. E.2).

The following preprocessing steps were done for the PDAC data before training:

- ST count (*X*_t_) normalization to 10000 counts per spot
- Log1p transformation per spot-gene pair across all spots and genes
- Standardization of UNI features (1024) for the histology (*X*_h_) data, per slide

The model was trained with a batch size of 256, for 50 epochs with the Adam (Kingma & Ba, 2014) optimizer at a learning rate of 0.001.

After training the model with all of the PDAC data, embeddings were extracted for the posterior distributions of the model: 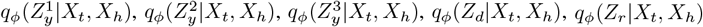. In this case, the means were not only used, and distributions for each posterior were sampled once per datapoint. For example, for 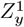 and sample *s* for spot *i* in the observed data:

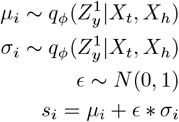

The samples for these posterior distributions were high dimensional (all 20 except for *Z*_d_ which was 5). Therefore, for visualization, these were reduced to 2-dimensions using PCA (Pedregosa et al., 2011), and the two axes of highest variation were visualized (Fig. 3 and 4).

The available cell-type labels derived from the ST data, pathologist annotations, and batch labels were overlaid on the posterior samples for each spot in the data, for the respective plots.

For this experiment, the SpatialDIVA model we trained had modified ELBO hyperparameters (*β*’s as defined in Appendix F). Specifically, *β*_1_ and *β*_2_, corresponding to the likelihood expectations, were set to 100. All other *β*’s remained 1.

#### H.2.2 Multi-modal batch correction benchmark

The batch-correction benchmark used a conditional variational autoencoder (cVAE) (Sohn et al., 2015) as a baseline, which conditioned on batch/slide label during training.

We trained SpatialDIVA and the cVAE using the exact same architecture and training setup as the quantitative disentanglement experiments (Sec. H.1.1). The key difference is that we performed this analysis for 5 iterations and in each iteration we used the entire dataset (colorectal or pancreatic cancer) for training and then subsequently evaluated batch-correction by freezing the network and getting embeddings for all datapoints. Training on the full dataset and obtaining batch-corrected embeddings is standard practice (Tran et al., 2020; Luecken et al., 2022).

Randomization in this case corresponded to different torch seeds for initialization of the models to capture training variability. The architecture of the cVAE model is the same as that of the VAE model in Sec. H.1.1, other than an added input of one-hot encoded batch-labels. This is done to reflect the supervision for batch-correction available to the SpatialDIVA model.

For evaluation, we extracted the 20 dimensional posterior means from the cVAE latent space 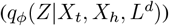, and the following 20 dimensional posterior means from SpatialDIVA: 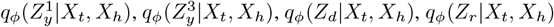.

The posterior means from each latent outlined from SpatialDIVA was used in the analysis, as well as the posterior means from the cVAE. We used the the scib-metrics package for benchmarking batch correction (Luecken et al., 2022). Specifically, we used the average silhouette width (ASW) (Luecken et al., 2022), which measures how well the batches mix across cells in given cell-types, and averages this value across cell-types. For each cell-type label *C*_j_ with *n* total cells:

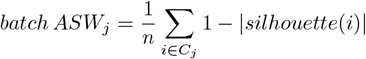

This value effectively measures how well the batches are mixed in the embeddings for a given cell-type *C*_j_, where 1 indicates perfect mixing and 0 indicates the most suboptimal mixing of batches possible. After obtaining this quantity per cell-type, we can average across *M* cell-types:

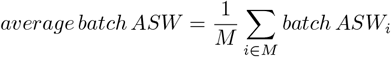

For cell-type labels, we combined the ST-celltypes and pathologist annotations for both the colorectal and pancreatic cancer datasets, into one string value which was then encoded for use with this metric. We calculated this metric for each of the latent subspaces indicated from SpatialDIVA and the latent subspace from cVAE. The calculation was done for 5 iterations for both cVAE and SpatialDIVA, as indicated.

#### H.2.3 Multi-modal biology conservation benchmark

For the results in Table 5, we used the exact same pipeline as in Sec. H.2.2, but instead of using the ASW metric, we used an aggregated score that measures how well the embeddings preserve biological signal with respect to the combined pathologist annotation and ST celltype labels. The metrics we used were also from the scib-metrics package (Luecken et al., 2022), and included the following bio-conservation metrics:

- Isolated biology label score
- Cell-type local inverse simpson index (cLISI)
- NMI using k-means clusters and biology labels
- ARI using k-means clusters and biology labels
- Biology label ASW

Full details on all of these metrics can be found in the scib-metrics documentation and the original scib publication (Luecken et al., 2022).

The values of these metrics for cVAE and SpatialDIVA were averaged per iteration to determine the biology conservation score.

### H.3 Results Sec. 4.4

#### H.3.1 Conditional multi-modal generation of PDAC

For this analysis, the PDAC data was processed using the following steps:

- Standardization of UNI features (1024) for the histology (*X*_h_) data, per slide

The model was trained on all of the genes, hence no highly-variable gene selection. Further, normalization and log transformation of the counts was not done as we considered a negative binomial likelihood for the ST data (*X*_t_), which works best with untransformed transcriptomic counts (Lopez et al., 2018).

The SpatialDIVA model was set up based on the changes to the default architecture as indicated in Sec. E.2. A negative binomial parametrization of the ST (*X*_t_) likelihood allowed for resampling of the counts after training the model, based on conditioning of certain covariates. We maximized a Negative Binomial likelihood, with the formulation taken from the scVI model (Lopez et al., 2018; Gayoso et al., 2022). The same formulation was used for subsequent sampling of Negative Binomial distributions using the obtained parameters.

The model was trained on all of the PDAC data, with a batch size of 256, the Adam optimizer (Kingma & Ba, 2014) at a learning rate of 0.001, and for 50 epochs.

After training, conditional multi-modal generation was done, as outlined in Sec. 4.3, after conditioning on transcriptomic 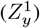, spatial 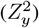 and morphological variation 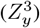, while holding other factors constant. This process was the same across all 3 covariates. As an example, for the intrinsic transcriptomic-conditioned generation for spot *i*:

##### Algorithm 1 Conditional generation of 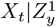

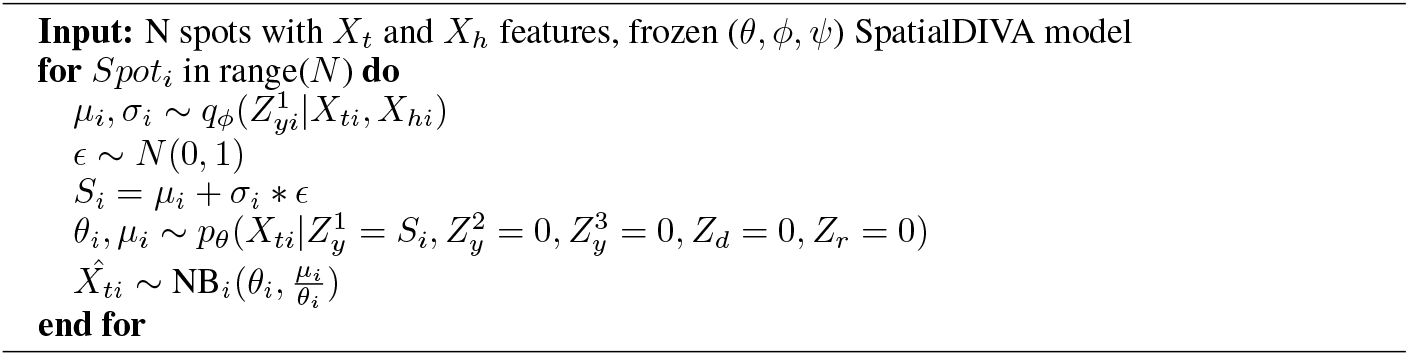

This conditional generation was done by conditioning on the three indicated covariates, for 10000 (*N*) randomly selected spots across the PDAC data. This resulted in three distributions of the transcriptomic counts, conditioned on each factor.

For each of these three distributions 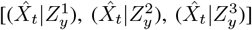, we performed the following processing steps to obtain the top 500 differentially expressed genes (DEGs) (Wolf et al., 2018):

- ST count 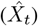 normalization to 10000 counts per spot
- Log1p transformation per spot-gene pair across all spots and genes
- Highly-variable gene selection using all 10000 spots (2500 genes)
- Count standardization and principal component reduction for the top 50 PCs
- Nearest-neighbor graph construction using the PCA embedding
- Leiden clustering of the data using the nearest-neighbor graph
- Differential gene expression across Leiden-derived clusters using the Wilcoxon rank-sum test

The scanpy (Wolf et al., 2018) library was used for these steps (v1.10.0). Default parameters were used, except where indicated. After determining the DEGs for each of the conditionally generated distributions, the top 500 were selected based on those exhibiting the highest Log-fold change in expression between clusters (Wolf et al., 2018). An all-versus one differential expression test was done, meaning that the expression in each cluster is compared with all of the other clusters to determine DEGs. Therefore, sorting by the highest Log-fold change can be interpreted as sorting genes that exhibit the highest specificity to their respective clusters in the reconstructed counts.

Using the top 500 DEGs from each of the conditionally generated count distributions 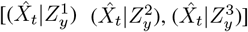, we used the gProfiler online platform (Kolberg et al., 2023) to perform pathway enrichment analysis. Default parameters were used, except for the following changes:

- ‘Ordered query’ was used
- The pathway databases were subset to GO molecular function, GO biological process, and REACTOME

Ordered query was utilized as the top 500 DEGs are ordered, and the subsetting of pathway databases was done to ensure that outdated and uncurated databases did not affect the analysis (Wadi et al., 2016).

The resulting enriched pathways were further analyzed for overlap. Pathways were sorted based on multiple-testing corrected *p*-values (Kolberg et al., 2023). The functional grouping of the pathways (Fig. 6a) was done based on presence of any of the following keywords:

- **Pancreas functions**: keywords = (“pancreas”, “pancreatic”, “development”, “glucagon”, “endocrine”, “metabolism”, “metabolic”, “catabolic”, “hormone”)
- **Immune functions**: keywords = (“immune”, “inflammatory”, “inflammation”, “cytokine”, “chemokine”, “interleukin”, “interferon”, “infection”, “immunoglobulin”, “antigen”)
- **Extracellular matrix functions**: keywords = (“extracellular”, “matrix”, “collagen”, “fibronectin”, “laminin”, “integrin”, “adhesion”, “cell-matrix”, “cell-cell”)

To determine the overlap between the top 500 DEGs and classical/basal subtype (Moffitt et al., 2015) markers, the following list of literature-derived markers were used:

**Basal**: - VGLL1 - S100A2 - LY6D - SPRR3 - SPRR1B - LEMD1 - KRT15 - CTSV - DHRS9 - AREG - CST6 - SERPINB3 - KRT6C - KRT6A - SERPINB4 - FAM83A - SCEL - FGFBP1 - KRT7 KRT17 - GPR87 - TNS4 - SLC2A1 - KRT5

**Classical**: - BTNL8 - FAM3D - PRR15L - AGR3 - CTSE - LINC00675 - LYZ - TFF2 - TFF1 - ANXA10 - LGALS4 - PLA2G10 - CEACAM6 - VSIG2 - TSPAN8 - ST6GALNAC1 - AGR2 - TFF3 CYP3A7 - MYO1A - CLRN3 - KRT20 - CDH17 - SPINK4 - REG4 - GATA6

## I Metrics to assess disentanglement in a continuous space

In this section, we provide a detailed explanation of the metrics outlined in Table 2. These metrics and the disentanglement evaluation framework are adapted from Carbonneau et al. (2020).

We define a set of *N* observations as *X* = **x**_1_, **x**_2_, …, **x**_N_. Each observation is assumed to be fully determined by a set of *M* factors 𝒱 = {*v*_1_, *v*_2_, …, *v*_M_} through a generative process *g*(**v**) ↦ **x**. Let *V* = {**v**_1_, **v**_2_, …, **v**_N_} represent the factor realizations that generate *X*. A representation learning algorithm maps *r*(**x**) ↦ **z**, where **z** ℝ^d^ is a point in the learned latent space Ƶ = {*z*_1_, *z*_2_, …, *z*_d_}. The set *Z* = {**z**_1_, **z**_2_, …, **z**_N_} contains all points in *X* projected into the latent space by *r*(.). The disentanglement metrics evaluate the relationship between *V* and *Z* to compute a disentanglement score.

In our setup, we consider *Z* to be the continuous learned latent space from embeddings of the baseline models and SpatialDIVA, and the one-hot encoded labels from the data (Appendix H) to be 𝒱.

The details of the metrics, originally outlined in Carbonneau et al. (2020), are indicated below. The terms latent space and latent codes are used interchangeably for *Z* = {**z**_1_, **z**_2_, …, **z**_N_}.

### I.1 Explicitness Score

As proposed in Ridgeway & Mozer (2018), explicitness is measured by training a classifier on the entire latent space to predict factor classes, assuming discrete factor values. A simple logistic regression classifier is employed, and its performance is evaluated using the area under the ROC curve (AUC-ROC). The final explicitness score is the average AUC-ROC across all classes and factors. Since the minimum AUC-ROC value is 0.5, the scores are normalized to fall within the range [0, 1]. The logistic regression loss is adjusted due to class imbalance.

### I.2 Attribute Predictability Score (SAP)

SAP (Kumar et al., 2018) assigns a score *S*_ij_ for every factor-code pair (*v*_i_, *z*_j_). For categorical factors, a decision tree classifier is used, and balanced accuracy is returned. Scores for codes below a user-defined energy threshold (*dead-codes*) are set to 0. The complete SAP score is calculated as the average difference between the two highest scores *S*_ij_ for each factor:

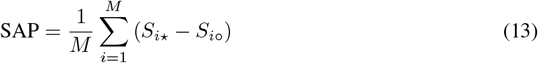

Here, *S*_i⋆_ and *S*_i°_ are the highest and second-highest scores for factor *v*_i_, respectively. Large differences indicate better disentanglement.

### I.3 Modularity Score

Modularity measures whether each code dimension *z*_j_ is associated with only one factor. Following Ridgeway & Mozer (2018), the factor *v*_⋆_ with the highest mutual information (MI) for each code dimension is identified, and the MI values with other factors are penalized:

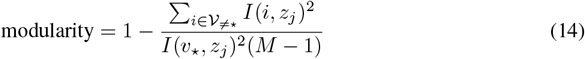

Here, 𝒱≠ _⋆_ is the set of all factors except *v*_⋆_, and *M* is the number of factors. The modularity score is averaged over all code dimensions.

### I.4 Mutual Information Gap (MIG) and MIG-SUP

MIG (Chen et al., 2018) evaluates the compactness of the representation by computing the MI between each factor and code dimension. The difference between the highest and second-highest MI for each factor is normalized by the factor entropy:

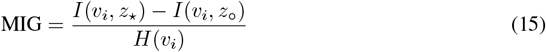

The MIG score is the average gap across all factors.

MIG-sup (Li et al., 2020) extends MIG to include modularity. It computes the MI gap from the perspective of the code:

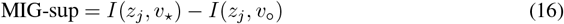

The meaningful code dimensions are determined by a threshold on *I*(*z*_j_, *v*_⋆_). All code dimensions are considered to avoid thresholding.

### I.5 Joint Entropy Minus Mutual Information Gap (JEMMIG)

JEMMIG (Do & Tran, 2020) addresses MIG’s inability to measure modularity by incorporating the joint entropy of the factor and its most related code dimension:

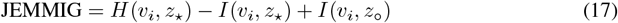

The score is normalized to lie between 0 and 1:

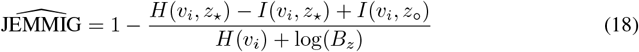

The average score across all factors is reported.

### I.6 DCI-MIG

DCIMIG (Sepliarskaia et al., 2020) combines elements of DCI and MIG. It computes MI gaps for each factor and code dimension and aggregates the scores into a single disentanglement measure:

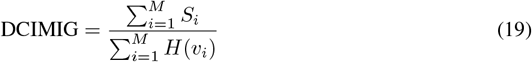

Here, *S*_i_ is derived from the maximum MI gap associated with each factor.

### I.7 Interventional Robustness Score (IRS)

IRS (Suter et al., 2019) quantifies the robustness of code dimensions to changes in nuisance factors. Sets of codes are compared before and after targeted interventions, and the maximum observed distances are used to compute the final score, weighted by factor realization frequencies.

## J Code and data availability

Code associated with preprocessing (Appendices C and D), creating the SpatialDIVA model (Appendix E), and running the experiments outlined in the paper (Appendix H) can be found at https://github.com/hsmaan/SpatialDIVA. The datasets for the CRC and PDAC cohorts will also be released and linked at the SpatialDIVA Github.

